# Use-dependent plasticity in human primary motor hand area: Synergistic interplay between training and immobilisation

**DOI:** 10.1101/217661

**Authors:** Estelle Raffin, Hartwig Roman Siebner

## Abstract

Training and immobilization are powerful drivers of use-dependent plasticity in human primary motor hand area (M1_HAND_). Here we used transcranial magnetic stimulation to clarify how training and immobilisation of a single finger interact within M1_HAND_. Healthy volunteers trained to track a moving target with a finger for one week. The tracking skill acquired with the trained finger was transferred to a non-trained finger of the same hand. The cortical representations of the trained and non-trained finger muscle converged in proportion with skill transfer. Finger immobilisation alone attenuated the corticomotor representation and pre-existing tracking skill of the immobilized finger. The detrimental effects of finger immobilization were blocked by concurrent training of the non-immobilized finger. Conversely, immobilization of the non-trained fingers accelerated learning during the first two days of training. The results provide novel insight into use-dependent cortical plasticity, revealing synergistic rather than competitive interaction patterns within M1_HAND_.

## Introduction

Use-dependent plasticity of motor representations in the primary motor hand area (M1_HAND_) plays a critical role for learning dexterous movements (Plautz et al., 2000; Mawase et al., 2017; Lemon, 1999). In humans, motor representations within M1_HAND_ are dynamically shaped by sensorimotor experience (Siebner and Rothwell, 2003; Classen et al., 1998). Use-dependent representational plasticity has been extensively studied in rodents (Alaverdashvili and Paterson, 2017; Kleim et al., 1998) and monkeys (Nudo and Milliken, 1996; Nudo et al., 1996; Schieber and Deuel, 1997), suggesting a competition between cortical motor representations In monkeys, trained representations in M1 expanded at the expense of the representational zones of the adjacent body parts (Nudo et al., 1996). In contrast, long-term sensorimotor immobilization led to shrinkage of the “restricted” corticomotor representations, boosting the adjacent representations as in monkeys and rodents (e.g. Milliken et al., 2013) (e.g. Viaro et al., 2014).

Plastic changes in corticomotor representations can be mapped non-invasively in human M1_HAND_ with Transcranial Magnetic Stimulation (TMS) (Thickbroom et al., 1999; Wassermann et al., 1992; Wilson et al., 1993; Kleim et al., 2007). Classically, a figure-of-eight shaped coil is discharged over a grid of scalp positions and the mean amplitude of the Motor Evoked Potentials (MEPs) is calculated for each grid site, enabling the construction of a corticomotor map for the target muscle. TMS-based corticomotor mapping revealed *use-dependent representational plasticity of single muscle representations* in M1_HAND_. Echoing the results obtained in animals, trained cortical muscle representations increased after repeated practice of simple or complex sequential movements (Classen et al., 1998; Muellbacher et al., 2001; Pascual-Leone et al., 1994), whereas forced immobilization attenuated corticomotor representation (Liepert et al., 1995). While these studies provided converging evidence that training and immobilization are powerful drivers for plasticity in M1_HAND_, it remains to be clarified how experience-driven changes of distinct motor representations within M1_HAND_ interact and determine within-area plasticity of human M1_HAND_.

To address this question, we investigated how finger-specific visuomotor training or immobilisation interactively shape representational plasticity within human M1_HAND_. We hypothesized that finger-specific training or finger-specific immobilization would impact on the skill level and cortical representation of the finger that was not targeted by the intervention (i.e., non-trained or non-immobilized finger).

Despite widespread and intermingled motor representations in primate M1_HAND_ (Georgopoulos et al., 1999), there is a consistent latero-medial somatotopic gradient of the abductor digiti minimi (ADM) and first dorsal interosseus (FDI) muscle (Beisteiner et al., 2004; Beisteiner et al., 2001; Gentner and Classen, 2006; Quandt et al., 2012). We have recently introduced a novel neuronavigated TMS mapping approach which readily reveals the somatotopic arrangement of the ADM and FDI representations within M1_HAND_ (Raffin et al., 2015) (Dubbioso et al., in prep.). Here we exploited this TMS mapping approach to probe within-area somatotopic re-arrangement of motor finger representations in response to training or immobilisation of specific fingers.

Our experimental approach enabled us to test whether within-area plasticity in M1_HAND_ is characterized by competition or cooperation. Training-induced strengthening of one motor representation may occur at the expense of the non-trained motor representations. This competition may be particularly expressed when one motor representation is strengthened by training and the other is weakened through immobilization. Alternatively, experience-induced plasticity induced by finger-specific changes in sensorimotor experience may be mutually synergistic, benefitting also motor representations that are not directly targeted by training. A cooperative and synergistic mode of interaction implies that training of one motor representation would not benefit from concurrently weakening another one by immobilization. The prediction would rather be that the strengthening of the trained motor representation would stabilize the deprived motor representation.

To test which mode of interaction characterizes within-area representational plasticity within human M1_HAND_, healthy right-handed volunteers performed two sessions of a visuomotor tracking task one week apart (Fig. 1a). The tracking task required subjects to tracking a moving line with the left index or little finger (Fig. 1a). The tracking task was programmed as application on a smartphone which was attached to a wooden platform. The wrist and the non-trained fingers were fixed to the platform with Velcro strap to stabilize their position and to minimize co-contraction during tracking.

**Figure 1A.**
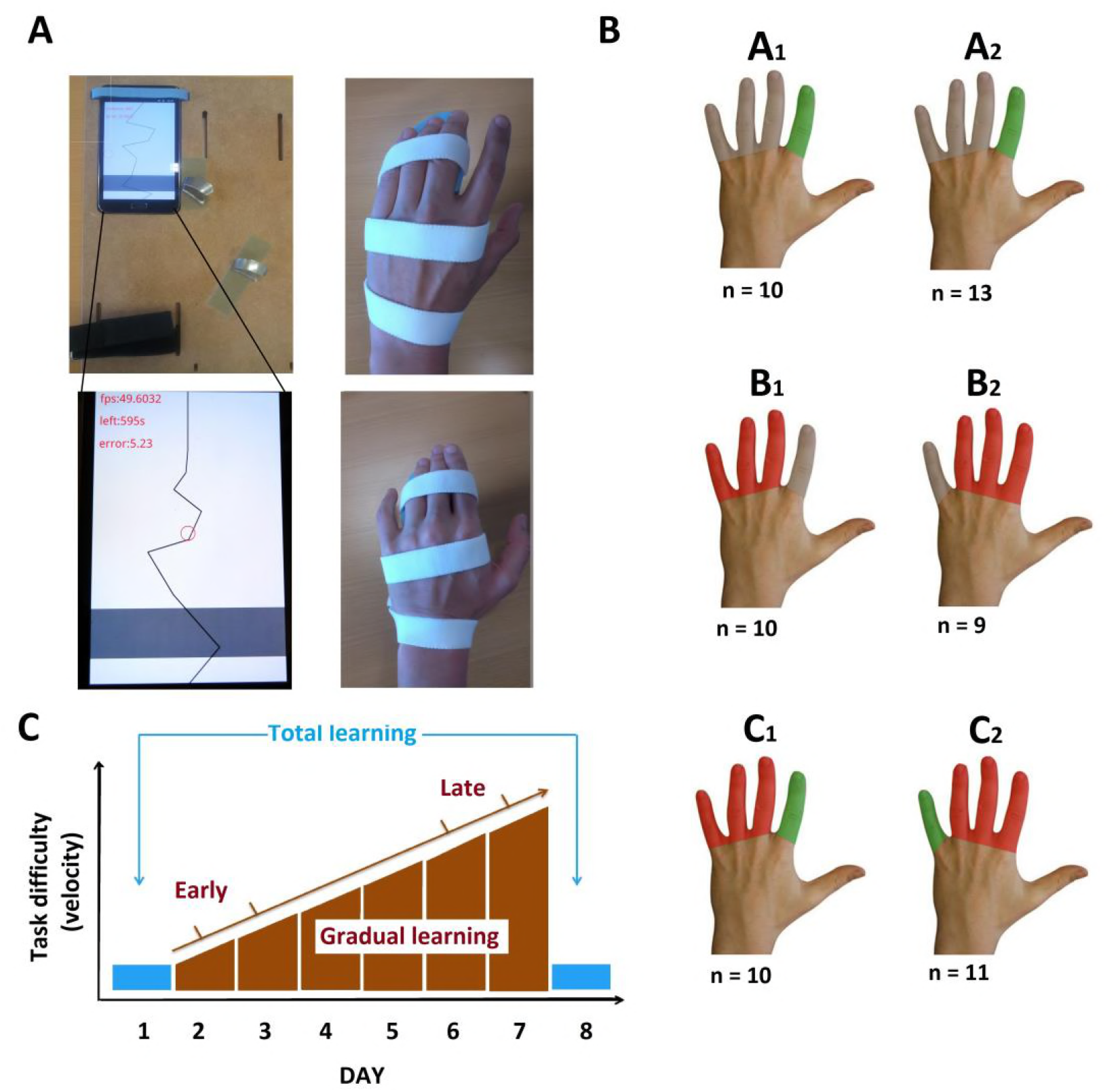
*Smartphone-based finger training using a flexible setup adjustable for training either the left index or little finger (left). The tracking task consisted in a moving line going from the top of the screen to the bottom. The red circle reflects the actual position of the subject’s training finger. This red circle was controlled by the index or index placed on the grey line. Feedbacks about the remaining time and online performances were provided. The right pictures display the immobilization procedure of three adjacent fingers (fingers III-V or II-IV) with an individually made splint; **1B:** Types of interventions: Group A1 and A2. Selective finger training without immobilization of adjacent fingers; Group B1 and B2: Immobilization of three adjacent fingers without training; Group C1 and C2: Selective finger training with simultaneous immobilization of adjacent fingers. Subgoups 1 and 2 differed in terms of the targeted finger; **1C:** Assessment of visuomotor tracking skill: Finger tracking with the index and little finger was assessed at day 1 and 8 using exactly the same task settings and during each training session at day 2 to 7 with a gradual increase in difficulty during consecutive sections.*

Participants were assigned to three groups, which were exposed to different sensorimotor experiences during the week between the two experimental sessions (Fig. 1b). Group A trained the same task with either their index or their little finger three times ten minutes a day, while task difficulty gradually increased from day to day. Group B underwent finger immobilization without any training (Group B). Group C received the same training as group A but with their adjacent fingers immobilized. Learning performances were quantified globally and gradually during the week using the absolute deviation between the target line and the movement performed by the subjects (Fig. 1c). Using neuronavigation, TMS was applied to seven M1_HAND_ targets which reflected the individual shape of the central sulcus (i.e., the “hand knob”) (Raffin et al., 2015). Sulcus-shape based TMS mapping was performed at baseline and after one week to capture experience-dependent changes in mediolateral cortical representations of left ADM and FDI muscles in the right M1_HAND_.

### Results

63 healthy volunteers were either exposed to one week of finger training, finger immobilization or finger training combined with immobilization of the remaining fingers. One week of finger-specific training or immobilisation was sufficient to shape dexterity as well as muscle-specific corticomotor representations in human M1_HAND_. Critically, each intervention had different effects on manual tracking skill and produced different patterns of within-area reorganization in human M1_HAND_.

### Changes in visuomotor tracking performance

We assessed the cumulative improvement in tracking ability using the percentage change in tracking accuracy at day 8 relative to baseline performance at day 1 (Fig. 2, left panel). Please note that the visuomotor tracking tasks performed at day 1 and 8 were matched in difficulty (Fig. 1c). A mixed ANOVA including all three interventional groups revealed a significant effect for the finger targeted by the interventions (F_(1,52)_ = 52.31, p < 0.001). This was due to an overall increase in tracking accuracy for the trained finger (Group A and C) or not immobilized (Group B) relative to the non-trained finger (Group A) or immobilized finger (Group B and C). The relative improvement in accuracy for the targeted finger depended on the type of intervention (F_(2,52)_ = 10.05, p < 0.001), while there was no systematic difference in the amount of overall learning between the little or index finger (F_(1,52)_ = 1.88, p = 0.18). A mixed ANOVA only including the data obtained in two learning groups (Group A and C) yielded similar results. There was a main effect for the *finger targeted by training* (F_(1,38)_ = 60.01, p < 0.001) and an interaction between *type of intervention* and *trained finger* (F_(1,38)_ = 33, p < 0.001).

**Figure 2.**
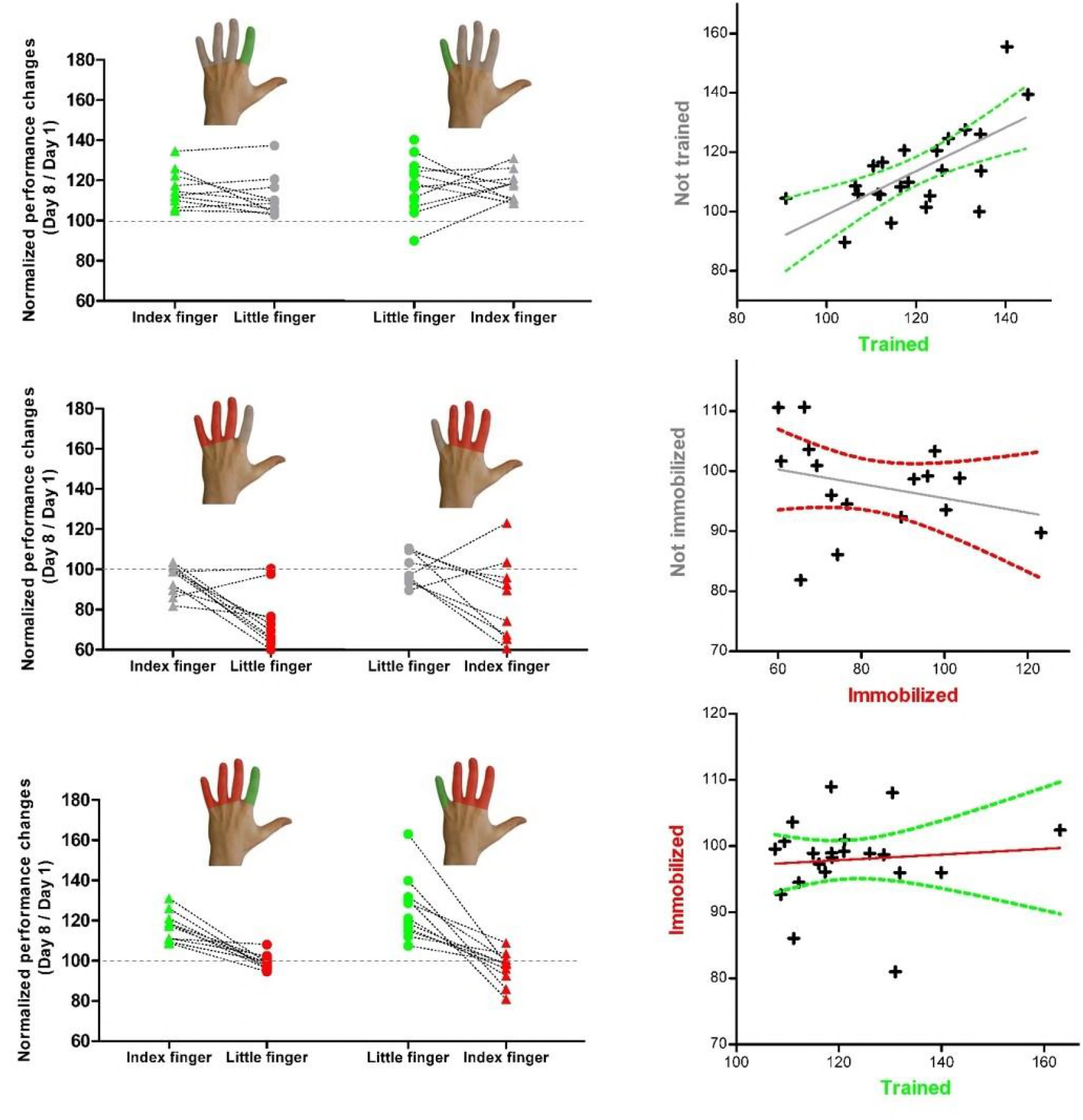
*Individual changes in tracking accuracy from day 1 to day 8. **Left panels.** The y-values reflect individual tracking accuracy at day 8 expressed as percentage of day 1 for each group. **Right panels.** The scatter graphs plot the individual performance changes for the two fingers of the same hand separately for each group. The straight grey line reflects the fit of the linear regression and the curved lines represent the 95% confidence interval.*

The significant interaction between the *type of intervention* and the *trained finger* motivated a follow-up analysis of overall learning within each interventional group. In group A, learning without concurrent immobilization only resulted in a trend advantage in tracking performance for the trained compared to the non-trained fingers (t_(22)_ = 1.94, p = 0.07). At the individual level, the improvement in tracking with the trained fingers correlated with improved tracking performance in the non-trained, non-immobilized finger (r = 0.66, p < 0.001; Fig. 2, upper right panel). In the “training only” group, the non-trained finger showed a significantly higher tracking accuracy at day 8 relative to the non-trained and non-immobilized finger in the “immobilization only” group (Group A vs group B, t_(40)_ = 4.85, p < 0.001). Together, the data indicate efficient transfer of the learned visuomotor tracking skill to the non-trained finger in the “training only” group (Fig. 2, upper panels).

In contrast, no learning transfer was found, when learning was combined with immobilisation (Group C). After one week of training, there were significant differences in tracking performances between the learned and the immobilized fingers (t_(20)_ = 7.88, p < 0.001) without any correlation among them (r = 0.1, p = 0.7; Fig. 2, lower panels).

Finger immobilization without concurrent training of the adjacent finger degraded visuomotor tracking ability of the immobilized finger (Group B, Fig. 2, middle panels). Pair-wise comparison showed a consistent decay in tracking performance at day 8 for the immobilized finger relative to the non-immobilized non-trained finger (t_(18)_ = 3.59, p = 0.002). The relative decrease in tracking accuracy in the immobilized finger did not correlate with tracking performance in the non-immobilized, non-trained finger (r = -0.28, p = 0.29), which showed similar tracking performance at day 1 and 8.

Concurrent immobilization of the non-trained fingers failed to boost the acquisition of the tracking skill in the trained finger. Tracking performance was comparable for the trained finger in group A and C (t_(42)_ = 1.14, p = 0.26), showing that overall learning was not enhanced by immobilization of the non-trained fingers in group C. However, concurrent training prevented degradation of tracking skill of the immobilized finger in group C (Fig. 2, lower left panel). The immobilized finger combined with training of the adjacent finger showed better tracking performance than participants in whom the finger was immobilized without concurrent training of the adjacent finger (Group C vs group B; t_(38)_ = 4.33, p < 0.001).

### Day-to-day changes in finger tracking performance

We analysed the behavioural data that had been recorded on the smartphone during home-based training sessions from day 2 to 7. Tracking accuracy was normalized to the gradual increase in difficulty level of the task from day to day. Daily training resulted in a gradual improvement of tracking skill (Fig. 3a). Mixed-effects ANOVA showed a main effect of *day of training* F_(3.24,37)_ = 15.6, p < 0.001) which did not differ between training with the index or little finger (F_(1,37)_ = 3.29, p = 0.08. While the total amount of performance improvement from baseline to day 8 was similar between group A and C, we found differences in the dynamics of day-to-day learning in the trained fingers between Group A and C (Fig. 3a & b). This was confirmed by a *day of training* by *type of intervention* interaction (F_(5,37)_ = 2.54, p = 0.03). The immobilization of the adjacent fingers accelerated early learning in group C. Group C showed a better tracking accuracy on days 3, 4 and 5 relative to Group A in which finger tracking was trained without concurrent immobilisation of the adjacent fingers (see Fig. 3a & b for the incremental learning curves for both trained fingers and Table 1 for post hoc t-tests comparisons).

**Table 1.** *Statistical results of post hoc t-tests comparing the normalized gradual learning across days in the two groups receiving training. Group A: Training without immobilization, group C: Training with immobilization of the adjacent fingers.*

| **Day** | **t values** | **Df** | **p values** |
| --- | --- | --- | --- |
| **2** | -1.29 | 39 | 0.21 |
| **3** | -3.27 | 39 | 0.002* |
| **4** | -3.35 | 39 | 0.002* |
| **5** | -2.65 | 39 | 0.012* |
| **6** | -1.35 | 39 | 0.19 |
| **7** | -1.84 | 39 | 0.07 |

**Figure 3.**
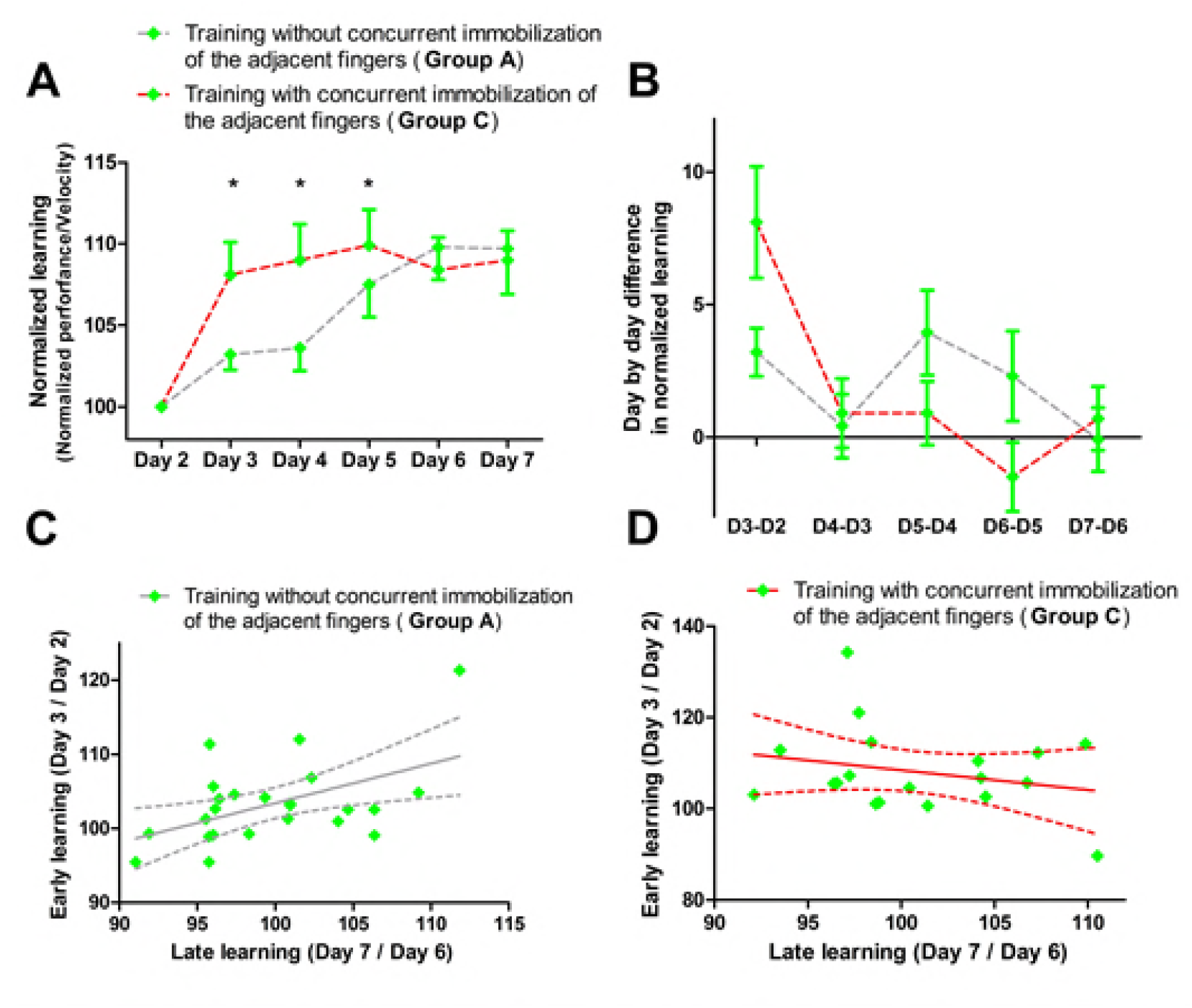
*Day-to-day improvement in tracking accuracy of the trained finger. Data from the index and little fingers are pooled together. For details regarding the calculation of the daily learning rate see the main text. **(A and B)** The panels show day-by-day improvements in visuomotor tracking for the trained fingers depending on the status of the adjacent fingers. Panel A shows the learning rate for each day. Panel B shows the difference in learning rate compared to the previous day. Training with immobilisation shows faster early learning than training without immobilisation. **(C and D)** The panels plot early day-to-day learning against late day-to-day learning for learning without (C) or with (D) immobilisation of the adjacent fingers. The straight grey line reflects the fit of the linear regression and the curved lines represent the 95% confidence interval. Early learning only scaled linearly with late learning when finger training was performed without concurrent immobilization of the adjacent fingers.*

When learning was performed without concurrent immobilisation, the amount of early learning (mean of day 2 and day 3) correlated with the magnitude of late learning (mean of day 6 and day 7), suggesting a linear increase in skill over consecutive days (Group A, r = 0.72, p < 0.001, Fig. 3c). This gradual continuous performance gain was not found when learning was combined with immobilisation of the adjacent fingers (Group C; r = -0.16, p = 0.49, Fig. 3d). Concurrent immobilization of the adjacent fingers modified the gradual build-up of skill from session to session during one week of training, acellerating early learning while flattening the slope of late learning. In group C, the rapid early increase in tracking performances (day 2 – day 3) scaled with the amount of cortical disinhibition in M1_HAND_ as reflected by the relative reduction in SICI from day 1 to day 8 (r = 0.54, p = 0.012, corrected for multiple comparisons).

### Experience-dependent within-area plasticity in right M1_HAND_

Sulcus-shape based TMS mapping was used to map the corticomotor representations of the left FDI and ADM muscles in each individual. Sulcus-shape based mapping showed that all interventions triggered a reorganization of cortical representations which involved changes in corticomotor excitability and spatial representation. (Fig. 4 & 5). Corticospinal excitability was measured as *Area Under the Curve* (AUC), representing the mean MEP amplitude for all seven-map positions. The ratio between AUC values obtained at day 8 (post-training) and day 1 (baseline) reflected relative changes in corticomotor excitability from day 1 to day 8.

**Figure 4.**
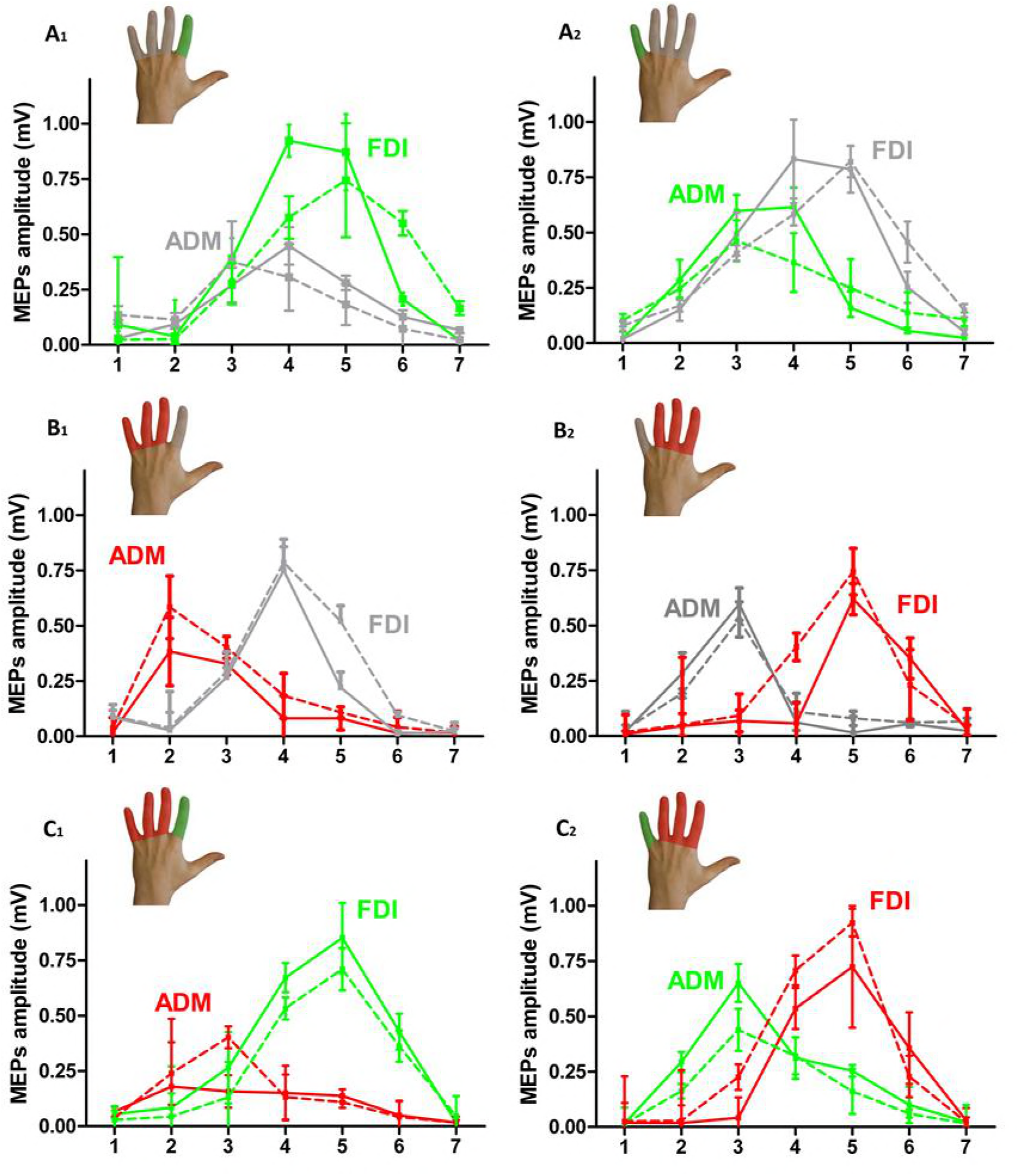
*Mediolateral cortical excitability profiles of the FDI and ADM muscle obtained with neuronavigated single-pulse TMS in the three experimental groups (A1/A2, B1/B2, C1/C2). The colour of the lines indicates whether the muscle was trained (green), immobilized (red), or neither immobilized nor trained (grey) on day 1 (dotted line) and day 8 (full line). Data points represent the mean value of each group. Error bars equal SEM.*

### Changes in regional corticospinal excitability

Visuomotor tracking training increased regional corticospinal excitability in the trained muscles regardless of which finger was trained (Fig. 4 & 5, panels A and C). Conversely, immobilisation alone attenuated corticospinal excitability of the immobilized muscle (Fig.4 & 5, panel B). The opposite effects of training and immobilization were reflected by a statistical interaction between *type of intervention* and *muscle* for the AUC ratio (F_(2,55)_=3.81, p=0.03). The bi-directional use-dependent change in corticospinal excitability did not differ between the FDI or ADM muscle (F_(1,55)_=0.16, p=0.69). There was also a main effect of *muscle* caused by larger AUC values for FDI relative to ADM muscle across all conditions (F_(1,55)_=40.63, p<0.001), presumably reflecting the higher relevance of the FDI muscle for dexterous movements during everyday life.

Follow-up comparisons revealed that one week of visuomotor finger training produced similar excitability increases in the training muscle regardless of whether the non-trained finger was immobilized or not (Group A vs. Group C: t_(42)_ = 0.75, p = 0.45). Immobilization only induced a reduction in AUC in the “immobilization only” group, but this reduction in corticospinal excitability of the immobilized muscle was prevented by concurrent training of the non-immobilized finger (group C vs. group B, t_(36)_=3,07, p = 0.004). Moreover, the “training-only” group showed larger AUCs of the non-trained finger muscle compared to the non-trained, non-immobilized finger muscle in the “immobilization only” group (group A vs. group B, t(38)=7,7, p < 0.001).

### Within-area reorganization in right M1_HAND_

Sulcus-shape based TMS mapping confirmed the well-known somatotopic arrangement of cortical finger representations in the M1_HAND_ with the FDI muscle being represented more laterally than the ADM muscle (Fig. 5). Accordingly, statistical comparison of mean MEP amplitudes at each stimulation position showed an interaction between *location of TMS* and *muscle* (F_(6,300)_ = 34.25, p < 0.001).

**Figure 5.**
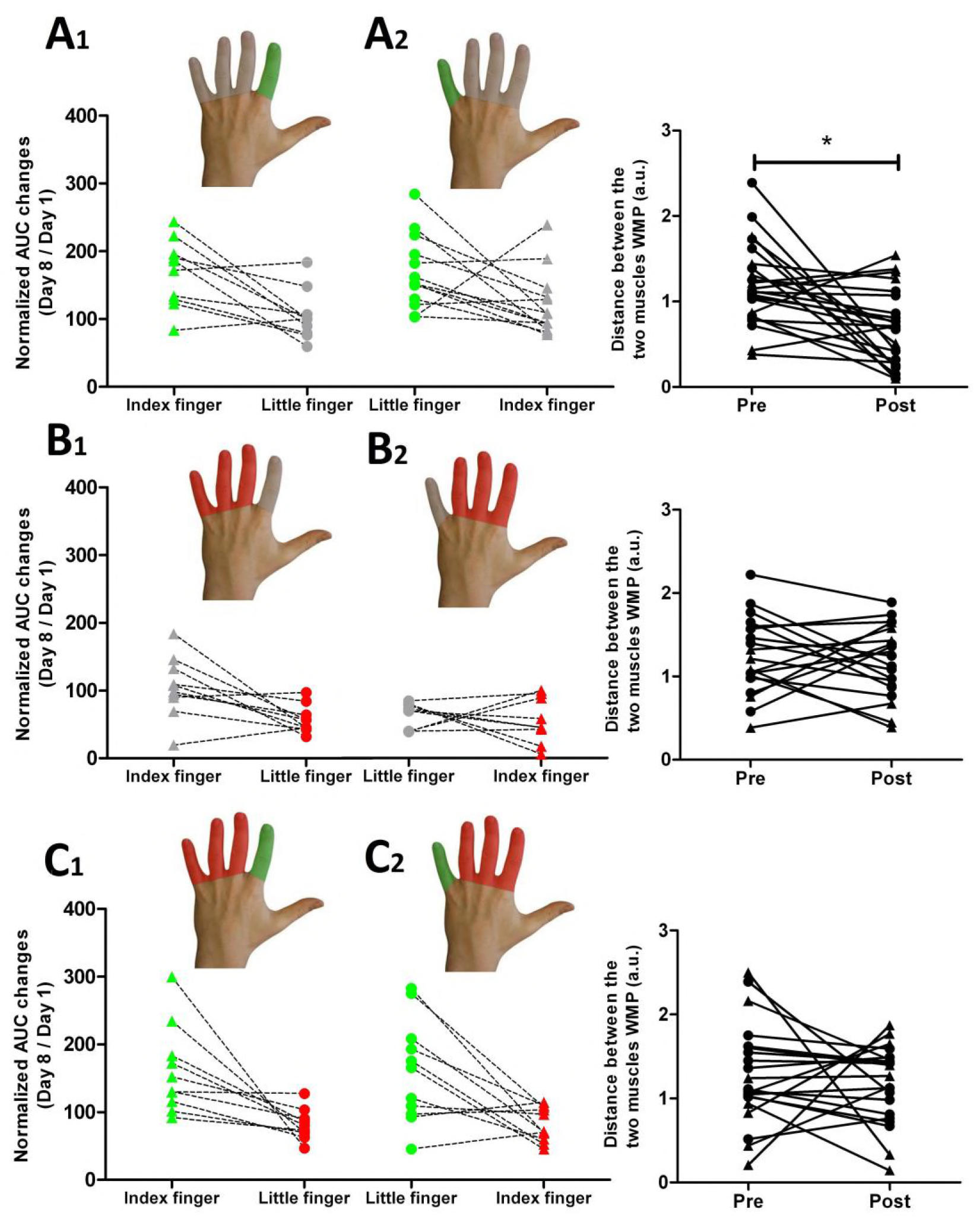
*Individual changes in mediolateral corticomotor representations of the left FDI and ADM muscles in right M1*_*HAND*_ *following finger-specific training or immobilization. Corticomotor representations were probed with sulcus-shape based single-pulse TMS mapping. **Left panels**: Relative changes in the area under the curve (AUC) from day 1 to day 8 given as percentage of baseline values. The colour of the lines indicates whether the muscle was trained (green), immobilized (red), or neither immobilized nor trained (grey); **Right panels:** Distance between the average mediolateral position of the muscle profiles (D*_*WMP*_*) before and after the intervention. Triangles symbolize the index finger and circles symbolize the little finger.*

Selective finger training resulted in a convergence of cortical muscle representations, but only when the non-trained fingers were mobile. The spatial representations of the FDI and ADM muscle in M1_HAND_ had moved towards each other after training, showing more overlap in group A, but not in group B and C. This pattern was confirmed by mixed-effects ANOVA which tested how the various interventions altered the distance between finger representations. We used the distance between the *Amplitude-Weighted Mean Position* (D_WMP_) of the FDI and ADM excitability profiles as index of spatial proximity between finger representations (see methods section). Mixed effects ANOVA revealed a change in spatial proximity between the FDI and ADM representation after one week relative to pre-interventional baseline (main effect of *session*: F_(1,57)_ = 6.7, p = 0.011). The spatial shift critically depended on the type of intervention, as indicated by an interaction between *session* and *type of intervention* (F_(2,55)_ = 3.32, p = 0.043). In the “training only” group, pairwise post-hoc t-tests showed that the mean position of the trained and non-trained muscle profiles shifted toward each other, resulting in smaller D_WMP_ values (group A; t_(22)_ = 3.45, p = 0.002, paired t-test). In contrast, mean D_WMP_ did not change in group B and C in which immobilisation was applied (p > 0.5).

### Experience-dependent changes in intracortical inhibition

Paired-pulse TMS mapping at an inter-stimulus interval of 2 ms was used to examine the magnitude or spatial distribution of short-latency intracortical inhibition (SICI). The overall strength of SICI, as reflected by the AUC of SICI across all stimulation sites (AUC_SICI_), was modified depending on the type of intervention. Only participants, who had been practicing visuomotor tracking movements for a week, showed reduced SICI in the trained muscle representation as revealed the mean AUC_SICI_ (Fig. 6). Mean AUC_SICI_ showed an interaction between *type of intervention* and *session* for SICI in the trained finger muscle (F_(2,56)_ = 1.4, p = 0.037). We calculated the ratio between AUC_SICI_ on day 8 and AUC_SICI_ at baseline to quantify the individual change in overall SICI. Using this variable, follow-up comparisons confirmed less SICI for the trained finger muscle representation in both training groups (Groups A and C) relative to the non-trained and non-immobilized muscle in group B which only underwent immobilization (Group A vs group B: t_(42)_ = 2.9, p = 0.006; Group C vs group B: t_(36)_ = 5.22, p < 0.001). No difference in AUC_SICI_ was found between the two training groups (Group A vs group C: t_(38)_ = 0.18, p = 0.86). While both training interventions reduced intracortical inhibition in the cortical representation of the trained muscle, they differed in terms of their impact on intracortical inhibition of the non-trained muscle representation. (Fig. 6). When finger training was not combined with immobilization, training-related disinhibition occurred in the cortical representations of both, the trained and non-trained muscles (Group A). In contrast, it remained restricted to the cortical representation of the trained muscle in individuals, in whom finger training was combined with immobilization (Group C). Considering only the two groups in which training was performed, ANOVA of SICI revealed an interaction between *type of intervention* and *muscle targeted by training* (F_(1,36)_ = 6.9, p = 0.012) and a main effect for the trained muscle (F_(1,36)_ = 24,96, p<0.001). Post-hoc analyses showed a difference between AUC_SICI_ of the trained and immobilized muscle in the group, in which training and immobilization were combined (Group C, t_(20)_ = 7.34, p < 0.0001). In contrast, there was no difference in AUC_SICI_ between the trained and non-trained muscle after training in the “training only” group (Group A, t(22) = 0.96, p = 0.35).

**Figure 6.**
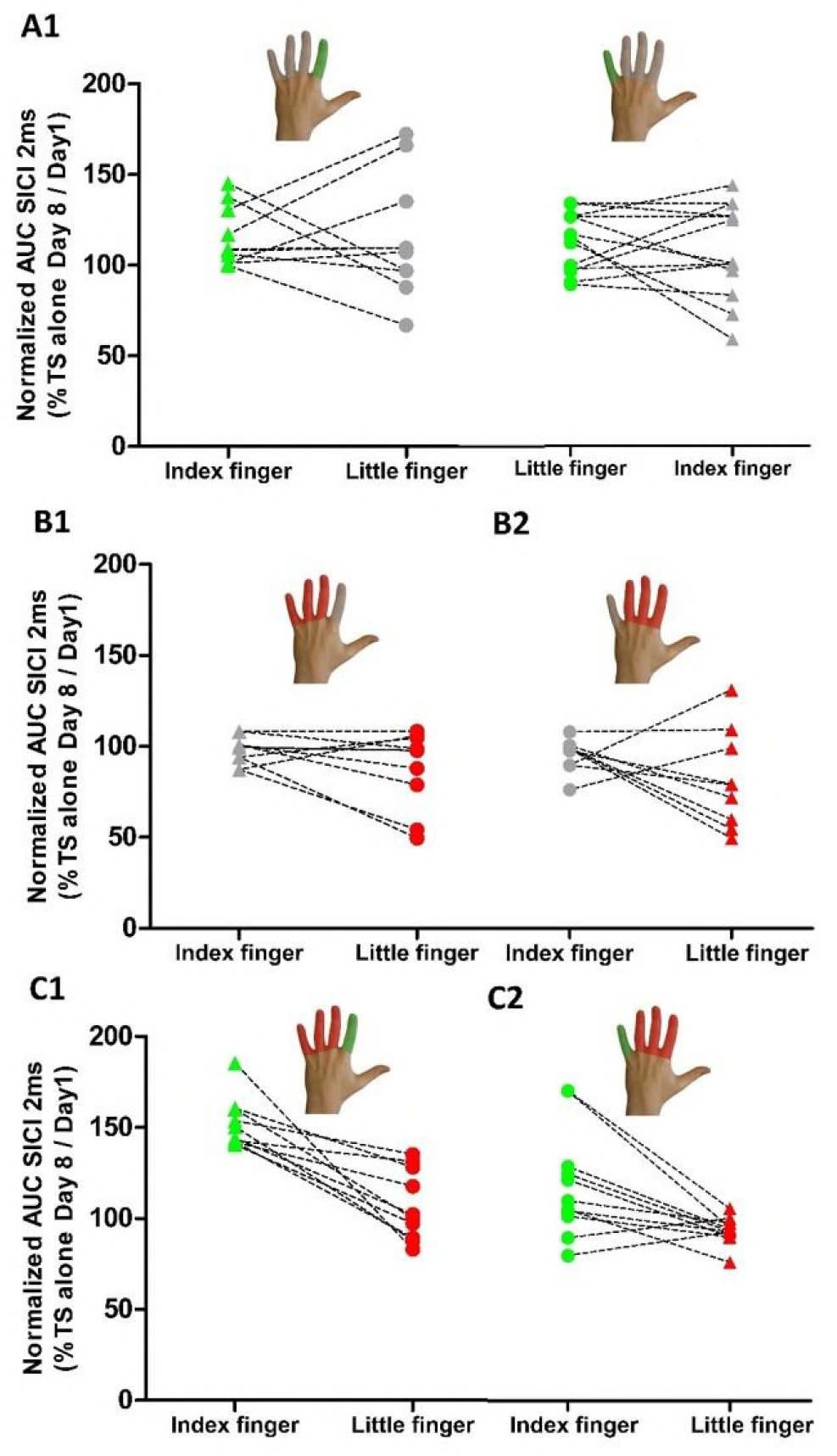
*Effects of finger-specific training or immobilization on mediolateral representations of short-latency intracortical inhibition (SICI) in M1*_*HAND*_ *probed with sulus-shape based, dual-pulse TMS.* The AUC_(SICI)_ at day 7 were expressed as percentage of AUC_(SICI)_ at baseline to capture relative changes in overall SICI after immobilization and training. ***Left panels.** Individual AUC_(SICI)_ ratios for the FDI and ADM muscle representations for the three types of interventions. An AUC ratio above the 100% line reflects a post-interventional decrease in SICI (i.e., disinhibition) relative to baseline. The colour of the lines indicates whether the muscle was trained (green), immobilized (red) or neither immobilized nor trained (grey); **Right panels.** Distances between the average mediolateral position of the SICI profiles (D_WMP_) are displayed before and after the intervention for the three main types of interventions. Triangles symbolize the index finger and circles symbolize the little finger.*

Immobilization alone increased intracortical inhibition in M1_HAND_. In the immobilized muscle, SICI increased from baseline to day 8 in individuals who underwent immobilisation without any training (Group B; Fig. 6). Immobilisation caused a relative decrease in AUC_(SICI)_ ratio, while the AUC_SICI_ ratio did not change in the non-immobilized, non-trained muscle, resulting in a significant difference between immobilized and non-immobilized muscle at day 8 (t_(18)_ = 2.33, p = 0.032).

In terms of spatial expression of SICI in M1_HAND_, the relative strength of SICI showed no clear difference in the relative magnitude of SICI among the cortical target sites. The spatial profile of conditioned MEP amplitudes followed those of the unconditioned MEPs evoked by the test pulse alone, showing that the relative magnitude of SICI was comparable across stimulation sites. Accordingly, ANOVA revealed no interaction between *location of TMS* and *Muscle* for SICI (F_(6,336)_ = 1.79, p = 0.1). None of the interventions had a consistent effect on the spatial arrangement of muscle-specific SICI profiles. Using the D_WMP_ values for the SICI excitability profiles as dependent variable, the mixed ANOVA revealed neither main effects nor interactions between *type of intervention* or *session* (p > 0.54).

### Relation between representational plasticity and visuomotor learning

We were interested to see whether our TMS-derived measures of representation plasticity would predict inter-individual differences in visuomotor skill learning of the trained finger or in learning transfer to the non-trained finger. To this end, we performed separate forward stepwise multiple regression analyses for the two training groups (Group A and C) treating the total learning scores as dependent variable. The D_WMP_ and AUC ratios of both finger muscles (FDI and ADM muscle) acquired with single-pulse and paired-pulse TMS were entered as potential predictors.

We first report the results regarding visuomotor learning of the trained finger. In the learning-only group (group A), the only TMS-based marker of representational plasticity that predicted the individual amount of training-induced visuomotor learning was the AUC increase of single-pulse MEPs in the trained muscle (Beta: 0.5, p=0.014; Table 2). The forward stepwise multiple regression model was significant (F_(1,21)_=7.14, p=0.014) and explained approximately 20% of the variance in overall finger tracking learning. For exploratory purposes, we also performed Pearson’s correlation analyses, which showed a positive correlation between learning from day 1 to day 8 and the relative AUC increase in the trained muscle (r = 0.5, p = 0.014, for all the other correlations: p> 0.05, corrected for multiple comparisons).

**Table 2:** *Regression analyses and predictive models for the learning transfer: Separate models were computed for group A and C. The following predictors were entered into the regressions as independent variables using a backward stepwise technique: total learning scores obtained by the adjacent finger, the distance of amplitude-weighted mean position* (*D*_*WMP*_) on the spTMS profiles, *and the area under the curve ratios acquired with single pulse and paired pulse TMS* (*AUC*_*SP*_ *and AUC*_*PP*_).

|  | Models |  |  |  | Significant predictors |  |  |
| --- | --- | --- | --- | --- | --- | --- | --- |
|  | Adj. R<sup>2</sup> | F-value | Df | P-value | Variable | Beta | P |
|  | Group A (training without immobilization of the adjacent fingers) |  |  |  |  |  |  |
|  | 0.22 | 7.14 | 21 | 0.014 | AUC<sub>sp</sub> | 0.5 | 0.014 |
|  | Group C (training with immobilization of the adjacent fingers) |  |  |  |  |  |  |
|  | Not significant |  |  |  | -- |  |  |
|  | Group A (training without immobilization of the adjacent fingers) |  |  |  |  |  |  |
|  | 0.23 | 7.48 | 21 | 0.012 | D<sub>WMP</sub> | -0.51 | 0.012 |
|  | Group C (training with immobilization of the adjacent fingers) |  |  |  |  |  |  |
|  | Not significant |  |  |  | -- |  |  |

In the group in which training and immobilization were combined (group C), the forward stepwise multiple regression model was not significant (*Table 2*). However, in line with the finding in group A, group C displayed a trend-wise positive correlation between the total learning and the AUC increase in the trained muscle (r = 0.42, p = 0.05).

We also tested which TMS-derived measure of representational plasticity predicts improvement in tracking skill in the non-trained muscle. In the learning-only group (group A), regression analysis revealed that the increasing proximity of the corticomotor representations of the FDI and ADM muscle predicted individual acquisition of visuomotor tracking skill with the non-trained finger (Beta: -0.51, p=0.012, *Table 2*). The forward stepwise multiple regression model on the total learning was significant (F_(1,21)_=7.48, p=0.012) and explained approximately 20% of total variance. The more the two muscle representations converged, the stronger was the amount of learning transfer to the non-trained muscle (r = -0.47, p = 0.023, for all the other correlations: p> 0.05). This was not the case in group C, the forward stepwise multiple regression model was non-significant (see *Table 2*), indicating that prevention of immobilization-induced skill degradation by concurrent training was not explained by any of the four TMS derived measures of representation plasticity.

## 5. Discussion

To the best of our knowledge, this is the first study showing that experience-induced representational plasticity of one motor representation can exert synergistic effects on another motor representation in human M1_HAND._ While there is an extensive behavioural literature demonstrating transfer of skill learning between hands (Laszlo et al., 1970; Schulze et al., 2002; Wang and Sainburg, 2004; Wang and Sainburg, 2004), this is the first demonstration that the motor system can transfer a learned visuomotor skill between single effectors of the hand (i.e. fingers). At the cortical level, learning transfer was paralleled by a convergence of finger muscle representations of the trained and non-trained finger in M1_HAND_ with the magnitude of convergence predicting skill transfer. By targeting the FDI or ADM muscle, we were able to internally replicate our findings. The three interventions induced analogous changes at the behavioural and representational level. This shows that the observed plasticity patterns can be generalized and were not specific for a given hand muscle.

Finger immobilization alone weakened the motor representation and impaired pre-existing tracking skill of the immobilized finger. Concurrent training with the non-immobilized finger neutralized the detrimental effects of finger immobilization. Conversely, immobilization of the non-trained fingers accelerated learning during the first two days of training without enhancing the total amount of skill improvement during the entire week of training. Figure 7 gives a synopsis of the reorganization patterns induced by the different types of interventions. In the following, we first discuss the cortical reorganization produced by visuomotor learning alone and then elaborate on how the learning-induced reorganization pattern was modified by concurrent immobilisation of the adjacent fingers.

**Figure 7.**
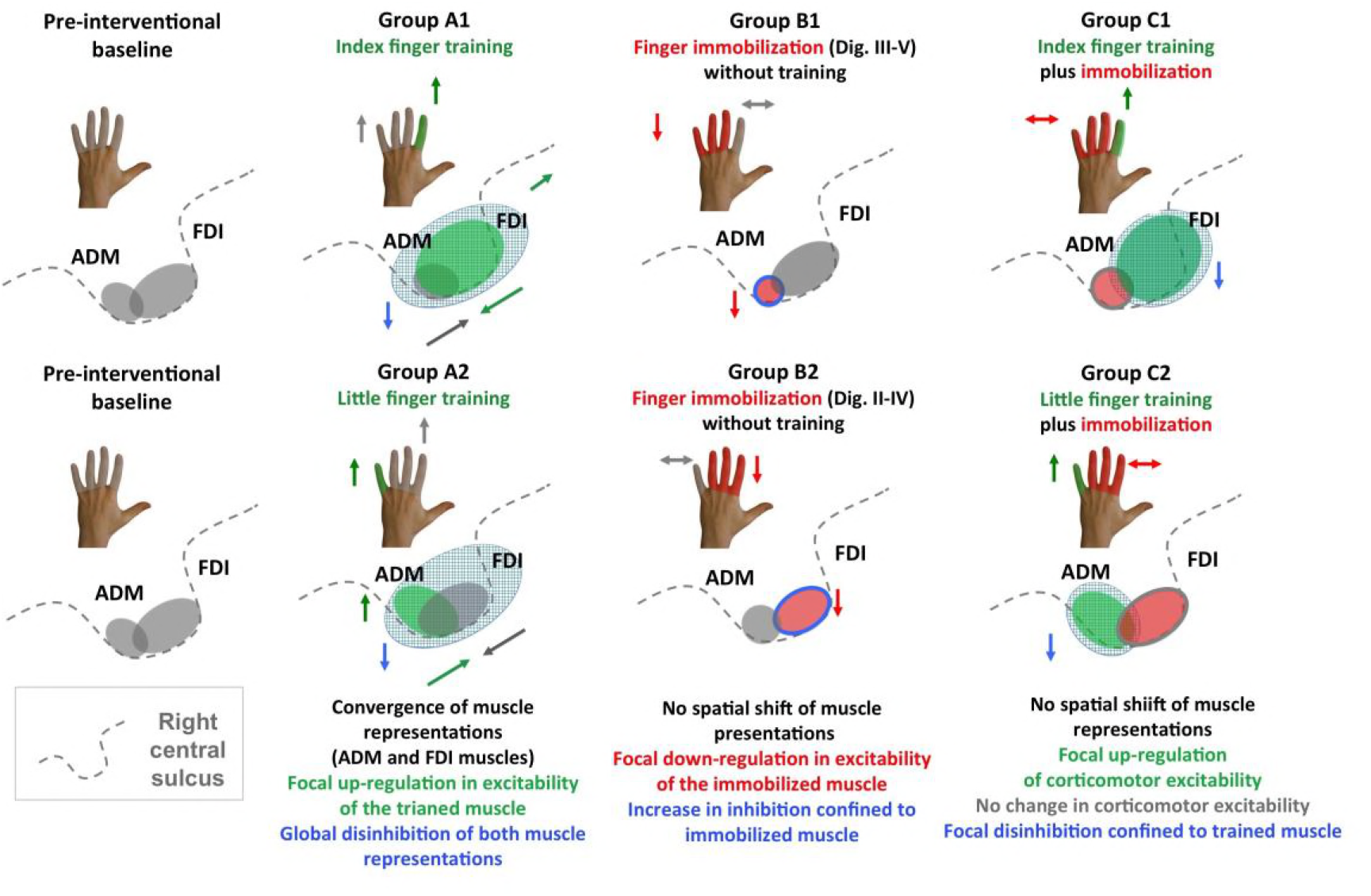
*Synopsis of within-area reorganization in right M1*_*HAND*_ *observed in Group A*_*1*_, *B*_*1*_, *C*_*1*_ and *A*_*2*_, *B*_*2*_ *and C*_*2*_. *The left panels illustrate the pre-interventional state with the grey areas reflecting the cortical representations of the FDI and ADM muscle in the right M1_HAND_. The arrows close to the schematic drawings of the hand summarize changes in learning performances for the trained and non-trained fingers. The grey shading illustrates “absence of intervention”, the green shading illustrates “training”, and the red shading illustrates “immobilization”. The arrows close to the schematic drawing of the central sulcus illustrate the direction of intervention-specific changes in muscle representations and intracortical inhibition.*

Training a visuomotor finger tracking skill shaped the corticomotor representation of the trained as well as the non-trained muscle (Fig. 7, group A1 and A2). One week of finger tracking training boosted the representation strength of the trained muscle representation, increased the spatial overlap, and attenuated intracortical inhibition of the trained and non-trained finger muscle of the same hand (Fig. 7). The overall increase in corticomotor excitability of the trained muscle predicted the individual amount of practice-induced visuomotor learning. This finding is in agreement with previous animal studies (Kleim et al., 1998; Nudo et al., 1996; Molina-Luna et al., 2008; Pruitt et al., 2016) or grid-based TMS mapping (Pascual-Leone et al., 1995; Svensson et al., 2003; Boudreau et al., 2013; Kleim et al., 2006; Tyc and Boyadjian, 2006) showing an expansion of the cortical representational maps of the trained body part. Likewise, there is consistent evidence showing that learning-induced up-scaling of corticomotor excitability in the trained muscle supports the acquisition of novel motor skills (Bagce et al., 2013; Koeneke et al., 2006).

In addition to an overall strengthening of the trained corticomotor representation, a spatial reorganisation within M1_HAND_ emerged over the course of one week (Fig. 7, group A1 and A2). Finger tracking training shortened the distance between the two mean positions of the trained and non-trained cortical motor representations. The convergence of corticomotor representations within M1_HAND_ predicted individual transfer of the learned tracking skill to the non-trained finger. The more the cortical representations converged, the higher the learning transfer to the non-trained muscle.

Using cortical microstimulation, previous animal studies showed a shift towards the motor territory of the adjacent non-trained body parts or an increased overlap with neighbouring representations of adjacent non-trained body parts (Kleim et al., 1998; Nudo et al., 1996; Molina-Luna et al., 2008). Our findings significantly extend these studies in two directions. Firstly, we show that a partial fusion of cortical motor representations does also occur within the cortical motor area presenting the same body part. Secondly, the results indicate that learning transfer of motor skills may at least partially be mediated within the primary motor cortex, possibly through a stronger overlap of functional representations. The prevailing notion is that learning transfer is mainly mediated through intermediate motor representations in premotor and parietal areas, which encode general knowledge of visuomotor predictions and skills (Grafton et al., 1998; Romei et al., 2009; Diedrichsen and Kornysheva, 2015). Our finding raises the possibility that some learning transfer might actually occur at the executive level in the M1_HAND_ through shared cortical motor representations. This hypothesis is in line with a recent study showing that the “trained” motor representation may contribute to intermanual transfer by “educating” the untrained motor representation or supporting the exchange of information between them (Gabitov et al., 2015).

Paired-pulse TMS of gamma-aminobutyric acid (GABA) mediated, intracortical inhibition revealed an attenuation of intracortical inhibition in contralateral M1_HAND_ after one week of training. Sulcus-shape based TMS mapping revealed that training-induced intracortical disinhibition was not confined to a distinct cortical site or to a specific muscle representation. On the contrary, the reduction in SICI was evenly expressed across all stimulation sites in M1_HAND_ and comprised the representation of the non-trained muscle. These observations significantly extend previous paired-pulse TMS studies which found training-induced reductions in intracortical inhibition (Stavrinos and Coxon, 2017; Coxon et al., 2014; Rosenkranz et al., 2007; Cirillo et al., 2011), showing that selective motor skill training with a single finger produces wide-spread disinhibition in M1_HAND_. Previous studies have shown that a reduction of gamma-aminobutyric acid (GABA) mediated, intracortical inhibition promotes synaptic plasticity in motor cortex and hereby, motor skill learning (Hess and Donoghue, 1994; Jacobs and Donoghue, 1991; Castro-Alamancos et al., 1995; Rioult-Pedotti et al., 1998). However, in the present study, the individual magnitude of SICI reduction did not scale with overall improvement in tracking performance after one week of training. The amount of disinhibition also did not predict the amount of skill transfer to the non-trained muscle. We therefore conclude that selective finger tracking training produces a widespread disinhibition of corticomotor representations in the “trained” M1_HAND_. Although GABAergic disinhibition, as measured with the SICI paradigm, may facilitate the expression of synaptic plasticity, it might not determine the final level of visuomotor tracking skill that can be acquired during one week of training. As we will discuss in more detail below, this might be different during early motor skill training, during which the focality and magnitude of intracortical inhibition might be more relevant. When the adjacent fingers were immobilized, selective finger training produced a more confined cortical reorganisation pattern (Fig.7, Group C1 and C2). Training enhanced the corticomotor representation of the trained muscle but not the non-trained, immobilized muscle without producing any spatial shifts. Like the increase in corticospinal excitability, training-induced cortical disinhibition was only expressed in the trained muscle. At the behavioural level, the magnitude of acquired tracking skill in the trained muscle was not enhanced after one-week of training as opposed to finger training alone. Training also produced no learning transfer to the non-trained muscle, when the non-trained muscle was immobilized. The effects of immobilization on training-induced plasticity and skill learning clearly speak against the notion that cortical motor representations are competing with each other for neural resources in the human M1_HAND_. If this were the case, immobilization-induced sensorimotor deprivation would have promoted an expansion of the trained muscle representation into the “deprived cortex” and hereby, boosted the learning success of the trained finger.

When training was combined with immobilization, sulcus-shape based TMS mapping of SICI revealed a more selective disinhibition of intracortical GABAergic circuits in the M1_HAND_ (Fig.7, Group B1 and B2). Relative reduction in SICI was limited to the trained muscle, while the immobilized muscle showed no consistent change (Fig. 8). We hypothesize that the muscle-specific attenuation of intracortical disinhibition in the trained muscle might have contributed to a faster learning rate during the first days of learning in the combined learning-immobilisation group. This hypothesis is supported by the observation that the rapid increase in tracking performances correlated with the reduction in SICI obtained after one week. Although speculative, it is possible that SICI reduction facilitates skill acquisition especially at the early phase of learning, while it functional role becomes less prominent during continued learning. This is in accordance with a recent study showing an early decrease in SICI after one day of learning and no change later on (Spampinato and Celnik, 2017). Furthermore, rapid GABAergic disinhibition can be induced acutely in M1_HAND_ by ischemic nerve block and has been shown to locally boost the expression long-term potentiation-like plasticity (Ziemann et al., 1998).

The modulatory influence of concurrent immobilisation of the adjacent fingers on training induced plasticity in M1_HAND_ can only be fully understood, when one considers the effects of immobilization alone on the corticomotor representations and visuomotor tracking skill (Fig. 7; group B1 and B2). Finger immobilization led to a selective down-regulation of corticomotor excitability with an increase in SICI, which was confined to the corticomotor representation of the immobilized muscle. The immobilized finger also showed a degradation of visuomotor tracking performance relative to pre-immobilization baseline. The findings indicate that one week of reduced sensorimotor experience is sufficient to weaken the deprived cortical representation and to deteriorate associated sensorimotor skills. These detrimental effects of finger immobilization were prevented by concurrent training of the non-immobilized fingers. Visuomotor tracking training of the neighbouring sensorimotor representation stabilized the pre-existing excitability and skill level of the immobilized muscle (Fig. 7). In line with previous animal data suggesting that recovery of a lesioned area depends on the activity of the adjacent cortical regions (Castro-Alamancos and Borrel, 1995), our findings provide additional support for a collaborative and synergistic mode of interaction between motor representations within M1_HAND_: The combined intervention resulted in a relative “up-scaling” of both muscle representations in M1_HAND_, increasing the trained muscle representation and preserving the immobilized muscle representation. Likewise, the net effect of finger training on dexterity was synergistic, improving the tracking skill in the trained muscle and maintaining the pre-existing skill level in the non-trained muscle despite of immobilization-induced deprivation.

Our findings have practical implications for preserving or recovering manual motor skills. In patients, in whom the upper limb has to be partially immobilized, intensive motor training of the non-immobilized part of the limb may help to minimize a functional degradation of motor skills relying on the immobilized muscles. Besides, immobilisation of the non-affected limb is a commonly used strategy to boost motor function of the affected limb in patients with chronic motor stroke (Taub et al., 1993; Taub and Morris, 2001; Taub and Uswatt, 2006; Morris et al., 1997). While constraint induced movement therapy may improve motor function of the affected limb, immobilization of the non-affected limb is likely to weaken the “immobilized” corticomotor representations in the healthy non-lesioned hemisphere. Future studies are warranted which systematically assess the effects of constraint induced movement therapy on the motor representations in the healthy non-lesioned hemisphere and how this might affect skilled hand function of the intact limb.

## Methods

### Participants

63 healthy individuals (25 females, age range: 19 – 48 years) participated in the study. Participants had no history of neurological or psychiatric illness and took no centrally acting drugs. Only individuals with little (less than 2 years) or no formal music training were included. All participant were strongly right handed according to the Edinburg Handedness Inventory (Oldfield, 1971). Prior to the study all participants gave written informed consent according to a protocol approved by the Ethical Committees of the Capital Region of Denmark (H-4-2012-106).

### Experimental design

Using a parallel-group design, participants were randomly assigned to one of three interventions (Fig. 1b). Group A (n=23, 12 females, mean age: 27.4 years) had to train a visuomotor tracking task for one week. The tracking task was programmed as application on a smartphone. The smartphone was attached to a wooden platform. The wrist and the non-trained fingers were fixed to the platform with Velcro strap to stabilize their position and to minimize co-contraction during tracking (Fig. 1a). At the inclusion (Day 0), we performed a careful multi-channel EMG measurement to ensure that participants were only activating the target muscle during tracking while keeping all other muscles relaxed.

Participants were required to make smooth abduction-adduction finger movements to follow a moving dot on the smartphone screen. Visuomotor tracking was either carried out with the left index finger involving the first dorsal interosseus (FDI) muscle (group A_1_; n= 10) or left little finger involving the abductor digiti minimi (ADM) muscle (group A_2_; n=13). Group B (n=19, 7 females; mean age: 26.1 years) performed no training, but digits III to V (Group B_1_; n= 10) or digits II to IV (Group B_2_; n=9) were immobilized. Group C (n=21, 8 females; mean age: 28.4 years) performed the same training task as group A for one week, but the adjacent fingers were concurrently immobilized. 10 participants (Group C_1_) trained with the index finger, while digits III to V were immobilized (Figure 1b). 11 participants (Group C_2_) trained with the little finger, while digits II to IV were immobilized. Visuomotor tracking performance was assessed in the laboratory at baseline (day 1) and post-intervention (day 8) using the same tracking task as for training. Performance was tested at a low difficulty level, which was identical for day 1 and 8 (level 1).

Using neuronavigation, sulcus-shape based TMS mapping of the corticomotor representations of the left FDI and ADM muscles was carried out on day 1 and 8. We applied single-pulse TMS to trace changes in the spatial profile of FDI and ADM representations along the hand knob in the right primary motor hand area (M1_HAND_). We performed the same mapping procedure with paired-pulse TMS to assess changes in magnitude and spatial distribution of short-latency intracortical inhibition (SICI). We also performed functional MRI during visuomotor tracking on day 1 and 8. These data will be reported separately.

### Finger tracking training

Participants assigned to group A or C performed daily visuomotor tracking exercises with a dedicated smartphone for one week (Fig.1c). Participants had to track a moving line with a dot controlled by their index or little finger. Daily training lasted 30 minutes and was distributed over three separate sessions to avoid fatigue. The difficulty of visuomotor tracking was step-wise increased from day 2 to day 7 and tracking performance was recorded on the smartphone. The velocity and the range of motion on the horizontal axis increased sequentially from level 1 (baseline level) to level 24 (highest level) to allow fair comparison between subjects. Hence, the tracking task became gradually more challenging for all the participants across the training week, starting from really slow movements requiring a maximum of 20 degrees of abduction-adduction to fast tracking requiring 60 degrees abduction-adduction. The time line of visuomotor training is illustrated in Figure 1d.

### Finger immobilization

In group B or C, three adjacent fingers were immobilized in a syndactily-like position for the entire week (day 1-7) by means of an individually shaped splint. The splint was made up of a rigid plastic form, covered with soft tissue, placed at the level of second phalangeal joint. We took care to ensure that the fingers were immobilized in a physiological position to prevent pain, swelling, or excessive sweating. The device was effective in restricting abduction–adduction and flexion–extension movements of the constrained fingers. Subjects were still able to perform a number of daily-life motor activities with the non-immobilized fingers of the left hand. Splint-wearing participants were only allowed to remove the splint during their daily washing procedures. In group C, participants performed additional training and were asked to take the splint off for training to match training conditions to group A (training without immobilisation). All participants tolerated immobilization without reporting problems. In particular, none of them experienced sustained pain during or after wearing the splint.

### Transcranial magnetic stimulation

#### Resting motor threshold

First, the site at which a single TMS pulse elicited a maximal motor response was determined for the left FDI muscle. The resting motor threshold (RMT_FDI_) was then determined at this stimulation site using the Parameter Estimation by Sequential Testing (MLS-PEST) approach (Awiszus, 2003). Stimulus intensity of TMS was adjusted to individual RMT of the FDI muscle (RMT_FDI_).

#### Sulcus-shape based, linear TMS mapping of M1_HAND_

We applied a novel linear mapping approach, which we have recently developed in our laboratory to study the somatotopic representation of the intrinsic hand muscles in human M1_HAND_ *(Raffin et al., 2015).* The mapping approach uses neuronavigation to deliver TMS at equidistant sites along a line that follows the individual shape of the central sulcus forming the so-called hand knob (Yousry et al., 1997). Our sulcus-shape based, linear TMS mapping method yields a one-dimensional spatial representation of the corticomuscular excitability profile in M1_HAND_ (Raffin et al., 2015). We stimulated seven targets placed along the bending of the right central sulcus with a coil orientation producing a tissue current perpendicular to the wall of the central sulcus at the target site. The order of target stimulation was varies across subjects but maintained constant within subjects. Each of the seven targets was first stimulated with 10 single TMS pulses followed by 10 paired TMS pulses. Single-pulse TMS was applied at an intensity of 120% RMT_FDI_.

Paired-pulse TMS was used to measure the magnitude and spatial distribution of short-interval intracortical inhibition (SICI) in M1_HAND_. Paired-pulse TMS used at an inter-stimulus interval of 2 ms. The intensity of the CS was set at 80% and the TS at 120% of RMT_FDI_ (Roshan et al., 2003). SICI is thought to be mainly mediated through gamma-aminobutyric acid-A (GABA-A) receptors (Ziemann et al., 1996). The magnitude of SICI is dynamically modified depending on the motor state. For example, SICI is reduced during voluntary contraction (Ridding and Rothwell, 1995; Opie et al., 2016) and is thought important for fractionated movement control (Zoghi et al., 2003). We performed paired-pulse TMS to trace changes in intracortical inhibition, because intracortical inhibition and cortical plasticity are tightly intertwined in M1_HAND_. A reduction of GABA-ergic intracortical inhibition has been shown to boost synaptic plasticity in motor cortex and to promote motor skill learning (Hess and Donoghue, 1994; Jacobs and Donoghue, 1991; Castro-Alamancos et al., 1995; Rioult-Pedotti et al., 1998). In humans, paired-pulse TMS measurements of SICI showed that motor training reduces intracortical inhibition which may contribute to training-induced plasticity (Stavrinos and Coxon, 2017; Coxon et al., 2014; Rosenkranz et al., 2007; Cirillo et al., 2011).

#### Electromyographic (EMG) recordings

Using a bipolar belly-tendon montage, motor evoked potentials (MEPs) were recorded with surface electrodes from the left abductor digiti minimi (ADM) and first dorsal interosseus (FDI) muscles during complete muscle relaxation (Ambu Neuroline 700, Ballerup, Copenhagen). The analogic signal was amplified and band-pass filtered (5-600 Hz) with a Digitimer eight-channel amplifier, digitized at a sampling rate of 5000 Hz using a 1201 micro Mk-II unit, and stored on a PC using Signal software (Cambridge Electronic Design, Cambridge, UK).

### Data analyses

#### Corticomotor mapping

Individual MEPs were visually inspected to remove trials with significant artefacts or EMG background activity (< 1%). The peak-to-peak amplitude of MEPs was extracted using Signal software in the time window between 10 and 40 ms after the TMS stimulus (Cambridge Electronic Design, Cambridge, UK). For the ADM and FDI muscle, we constructed medio-lateral corticomotor excitability profiles based on the mean MEP amplitudes for each TMS target site along the central sulcus forming the hand knob. We compared the medio-lateral distribution of mean MEP amplitudes in a mixed ANOVA, with the mean *MEP amplitude* evoked by single-pulse TMS at a given stimulation site as dependent variable. The *type of intervention* (group A vs. group B vs. group C) and which *finger received training or immobilization* (subgroup 1 [A_1_, B_1_ or C_1_] vs. subgroup 2 [A_2_, B_2_ or C_2_] were included as between-subject factors, while the *location of TMS* (target 1 to 7) and *session* (Day 1 vs. Day 8) and *muscle* (ADM vs. FDI) as within-subject factors.

We derived two complementary measures from the MEP-amplitude profiles to study in more detail dynamic changes in the muscle-specific representations in M1_HAND_. The *Area Under the Curve* (AUC) was taken as index sensitive to a global up- or down-scaling in corticomotor excitability. The *distance between* the *Amplitude-Weighted Mean Position* (WMP) of the FDI and ADM excitability profiles was used to assess changes in spatial proximity of muscle-specific corticomotor representations. The amplitude-WMP was calculated according to the following formula:

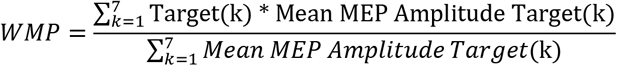

The AUC ratio (AUC at day 8/ AUC at day 1) and the distance between the WMP of the ADM and FDI muscle representation were analysed in separate mixed ANOVA models with *type of intervention* (group A, B, and C) and which *finger received training or immobilization* (subgroup 1 and 2) as between-subject factor. *Muscle* (FDI vs ADM) was added as additional within-subject factor to the ANOVA assessing AUC ratio. The factor *session* (day 1 and day 8) was only implemented in the ANOVA modelling WMP. The same statistical analysis was applied to the MEP-amplitude profiles evoked by paired-pulse stimulation at 2 ms using the normalized MEPs (Conditionned/Unconditionned MEPs) as dependent variable.

#### Relation between representational plasticity and visuomotor learning

We were interested to examine whether changes in AUC or in WMP distance obtained with single- or paired-pulse TMS would predict inter-individual differences in visuomotor skill learning of the trained finger in group A (training without immobilisation) and group C (training and immobilisation of adjacent fingers). To this end, we performed group-specific multiple regression analyses, treating the improvement in tracking performance of the trained finger from day 1 to day 8 as dependent variable. The predictive value of four TMS-derived measures were tested in a stepwise multiple regression model: (1) training-associated change in AUC assessed with single-pulse TMS (2) training-associated increase in AUC assessed with paired-pulse TMS at an ISI of 2 ms (3) training-associated change in distance between WMPs of the FDI and ADM excitability profiles assessed with single-pulse TMS (4) training-associated change in distance between WMPs of the FDI and ADM excitability profiles assessed with paired-pulse TMS at an ISI of 2 ms.

We conducted two additional regression analyses to examine whether changes in AUC or in WMP distance obtained with single- or paired-pulse TMS would predict the learning transfer of a visuomotor tracking skill from the trained to the non-trained finger in group A (training without immobilisation) and group B (training and immobilisation of adjacent fingers). We used the same stepwise multiple regression approach as described above. The only difference was that the total improvement in tracking performance of the non-trained finger from day 1 to day 8 as dependent variable.

Finally, another set of correlational analyses explored whether the changes in AUC or distance in WMP from day 1 to day 8 correlated with the amount of incremental learning (early learning score: Day 3/Day 2 and late learning score: Day 7/Day 6) and total learning (total learning score: Day 8/Day 1).

#### Visuomotor tracking

Learning of visuomotor tracking movements was assessed from two perspectives. To quantify the total amount of learning after the week of training (referred to as *total learning*), we compared the final tracking performance on day 8 with performance at baseline using a tracking task with the same difficulty level. To assess the gradual day-to-day improvement in tracking skill (referred to as *gradual learning*), we quantified mean tracking performance at each day of training and normalized this performance to the associated task velocity to take into account the increase in task difficulty.

Visuomotor performance was quantified using the mean relative error for each block, calculated as the difference in displacement between the tracking finger and a 3 mm target area centred around the target line every 100 ms of the task using a custom-made python script which calculated the percentage of time spent in the tolerance interval for each tracking block (in %).

The amount of *total learning* was determined by dividing the tracking performance measured on day 8 by initial performance on day 1 and expressed in percentage of improvement relative to baseline. We performed a global mixed ANOVA in which the improvement in tracking performance was treated as dependent variable. The *finger* (index vs. little finger) was treated as within-subject factor and the *type of intervention* (group A vs. group B vs. group C) and which *finger received training or immobilization* (i.e. subgroup 1 [A_1_, B_1_ and C_1_] vs. subgroup 2 [A_2_, B_2_ and C_2_] as between-subject factors. We also computed a more restricted ANOVA model which only included the groups that actually trained for one week, treating *finger* (index vs. little finger) as within-subject factor and the *type of intervention* (group A vs. group C) and which *finger received training* (i.e. subgroup 1 [A_1_ and C_1_] vs. subgroup 2 [A_2_ and C_2_] as between-subject factors. Conditional on significant main effects or interactions, we performed follow-up t-tests. In Group A and C, we tested whether total learning in the trained finger would predict the transfer of learning to the non-trained finger, using Pearson´s correlation.

To analyse *gradual learning* in group A and C, we multiplied the performance of a given training day with the corresponding tracking velocity to take into account the manipulation in task difficulty and normalized to the first training day. This measure was entered into a mixed effects ANOVA model with *day of training* (day 2 to day 7) and the *type of intervention* (group A and C) and which *finger received training* (subgroup 1 [A_1_ or C_1_] vs. subgroup 2 [A_2_ or C_2_]). We further tested for a correlation between early gradual learning (day 3 / day 2) and late gradual learning (day 7/ day 6).

#### Statistical considerations

All statistical analyses were performed using SPSS 19 for Windows (IBM, Armonk, NY, USA). The level of significance was defined as α = 0.05. Bonferroni-Sidak’s procedure was used to correct for multiple comparisons. Data are given as mean ± standard error of the mean (SEM). Normal distribution of the data was confirmed using the Kolmogorov–Smirnov test for all variables. For ANOVA, the Mauchly’s test of sphericity was performed. Greenhouse-Geisser correction method was applied to correct for non-sphericity.

## Acknowledgements

We would like to thank Arkadiusz Stopczynski for coding the smartphone application and all the volunteers for their participation.

